# Effects of a blend of essential oils in milk replacer on performance, rumen fermentation, blood parameters and health scores of dairy heifers

**DOI:** 10.1101/2020.03.17.995209

**Authors:** Joana P. Campolina, Sandra G. Coelho, Anna Luiza Belli, Fernanda S. Machado, Luiz G. R. Pereira, Thierry R. Tomich, Wanessa A. Carvalho, Rodrigo O. S. Silva, Alessandra L. Voorsluys, David V. Jacob, Mariana M. Campos

## Abstract

The objective of this study was to evaluate how the inclusion of a blend of essential oils in milk replacer (MR) affects different outcomes of dairy heifers. The outcomes evaluated: feed intake, performance, body development, blood cells and metabolites, insulin-like growth factor-1 (IGF-1), rumen fermentation, fecal scores and respiratory scores. All outcomes were evaluated during pre-weaning (4 – 60 d of age), and carryover effects on post-weaning (61 – 90 d of age) periods. The experimental units utilized were 29 newborn Holstein × Gyr crossbred dairy heifers, with genetic composition of 5/8 or more Holstein and 3/8 or less Gyr and body weight (BW) at birth of 32.2 ± 5.2 kg. Experimental units were randomly assigned to either a control group (CON, n = 15) or a treatment group, consisting of supplementation of a blend of essential oils (BEO, n = 14) with1 g/d/calf (Apex Calf, Adisseo, China). During the pre-weaning phase, all heifers were fed 5 L of MR/d reconstituted at 15% (dry matter basis), divided into two equal meals. Water and starter were provided *ad libitum*. During the post-weaning, animals received a maximum 3 kg of starter/d, and *ad libitum* corn silage, divided into two meals. The outcomes feed intake, fecal and respiratory scores were evaluated daily. BW was measured every three days, while body development was recorded weekly. Blood samples were collected on 0, 30 and 60 d of age for total blood cell count, weekly to determinate ß-hydroxybutyrate, urea and glucose, and biweekly for IGF-1. Ruminal parameters (pH, volatile fatty acids, ammonia-N and acetate:proprionate proportion - C2:C3) were measured each 14 days. A randomized complete block design with an interaction between treatment and week was the experimental method of choice to test the hypothesis of effect of BEO on all outcomes. An ANOVA procedure was used for continuous outcomes and a non-parametric test was used for the ordered categorical outcomes both adopting a C.I. = 95%. Results indicated that there was not enough evidence to accept the alternative hypothesis of effect of BEO in MR on: feed intake, performance, body development and blood metabolites during both pre-weaning and post-weaning periods. However, results indicated that the inclusion of BEO in MR significantly affects the proportion of C2:C3 during pre and post-weaning (P ≤ 0.05). Similarly, the effect is significant for basophil (P ≤ 0.001) and platelet (P ≤ 0.05) counts in pre-weaning. The interaction between week and treatment was also significant for lymphocytes (P ≤ 0.001), revealing a long-term treatment immunological effect. Lastly, the effect on fecal scores was also significant (P ≤ 0.05) during pre-weaning, with lower values for BEO. BEO contributed for ruminal manipulation on pre-weaning and carry over effect on post-weaning; immunity improvement and a decrease morbidity of neonatal diarrhea in pre-weaning phase.

## Introduction

A good calf-rearing program should embrace aspects that go from body development, stress reduction, good feed and housing management to optimum health status. Average daily gain (ADG) and body weight (BW) at weaning are important metrics used to measure the success of the rearing program. It is well known that these parameters are related to the heifer’s future milk production, as well as to its immune responses. Therefore, a bad life start can impact negatively on animal’s adult performance (1). Neonatal diseases, especially diarrhea, and nutritional mistakes can act as stressors, lowering calf immunity, and increasing animal susceptibility to other disorders (2).

The changes in intestinal and ruminal microbiota, caused by disease or malnutrition, can influence animal health, mortality rates and performance (3). Therefore, tools that help improving heifer development and health are essential to reduce disease, mortality and morbidity as well as accelerate the calf development. Additionally, since a calf is born functionally non-ruminant, it must have its digestive system and other organs and tissues changed in several weeks as well to adapt to the colonization of specific microbiota. The bacteria in the rumen must start the fermentation of carbohydrates, so the calf can become dependent mostly on volatile fatty acids (VFA) and not on lactose-driven metabolism anymore (4).

Since diseases have a negative impact on the heifer’s future performance, procedures that reduce the animal’s susceptibility to pathogens and stressors may improve future performance and productivity. Thus feed additives, such as growth promoter antibiotics, have been used to improve both rumen development and animal health (5), as well as increase performance and feed efficiency (6). However, the use of antibiotics in animal production for this purpose has been under severe criticism. Due to the increase of bacterial resistance and impact over human health it has been banned in several countries. Furthermore, the increase of organic dairy farms in the past years revealed a change in milk consumers’ preference with increased demand for “natural” and healthier foods. An alternative to the use of these additives is the utilization of herbal products, such as natural additives, for ruminant production (7).

Essential oils are herbal plant chemical components, constituted by volatile or ethereal oils that have been applied as a natural and safe alternative for antibiotics (8). Some of their properties are antiseptic and antimicrobial activities, that interfere on bacterial, fungal and protozoa cell functioning (9), presenting the similar efficiency to treat some diseases as antibiotics (10). They also contribute for prevention of oxidative stress (11), and help immune response to change leucocyte phagocytic activity and inhibiting complement system (12). Lastly, essential oils have a function similar to ionophores, influencing the gastrointestinal tract development, rumen microbiological activity, improving feed efficiency and decreasing neonatal diseases (9,13).

Previous studies have shown that essential oil supplementation in calf solid starter improves performance (14,15), rumen fermentation (16) and diarrhea severity (17). However, the effects over liquid diet supplementation are scarce. The aim of this study was to evaluate how adding a commercial blend of essential oils (BEO) in milk replacer (MR) affects feed intake, performance, feed efficiency, body development, blood cells and metabolites, insulin-like growth factor-1 (IGF-1), ruminal parameters, fecal and respiratory scores of dairy heifers during pre-weaning and post-weaning periods. Our hypothesis was that BEO supplementation in MR during pre-weaning would improve performance, influence blood parameters and health of dairy heifers.

## Material and methods

Protocols for this study were approved by the Ethics Committee of Embrapa Dairy Cattle (protocol number 9078250118). The experiment was conducted on the Embrapa Dairy Cattle Experimental Farm, located in Coronel Pacheco, Minas Gerais, Brazil, from March to September 2018.

### Animals, treatments and management

Twenty-nine newborn Holstein × Gyr crossbred dairy heifers, with genetic composition of 5/8 or more Holstein and 3/8 or less Gyr and BW at birth of 32.2 ± 5.2 kg, were used and equally distributed among treatments. They were separated from their dams immediately after birth and moved to individual sand-bedded pens (1.25 × 1.75 m, tethered with 1.2 m long chains), allocated in a barn with open sides and end-walls.

All heifers received 10% of their BW of good quality colostrum (Brix > 23%) up to 6 hours after birth, and had their umbilical cord immersed in iodine solution (10%).

At 2 to 3 d of age, heifers were fed 5 L/d of transition milk divided into two equal meals offered at 0800 and 1600 h, in buckets provided with rubber teats (Milkbar, New Zealand). At 3 d of age, blood samples were collected via jugular venipuncture with a clot activator tube (Labor Import, Osasco, Brazil), centrifuged at 1,800 × g for 10 min at room temperature (22 – 25 °C), to measure total serum protein using a Brix refractometer (Aichose refractometer, Xindacheng, Shandong, China). Heifers with low serum protein (Brix < 8.4%) were not enrolled in the present study.

Water and commercial calf starter (Soymax Rumen pre-inicial Flocculated, Total Alimentos, Três Corações, Brazil, Table 1) were offered in buckets for ad libitum intake (10% orts of solid feed).

**Table 1.** Nutrient composition (DM basis, % unless otherwise noted), of milk replacer (MR), starter and corn silage^1^.

| Item | MR<sup>2</sup> | Starter<sup>3</sup> | Corn Silage |
| --- | --- | --- | --- |
| DM (%) | 96.0 | 86.7 | 36.1 |
| CP (% of DM) | 19.4 | 17.1 | 7.9 |
| Ether extract (% of DM) | 14.1 | 3.9 | 4.3 |
| Organic Matter (% of DM) | 9.7 | 7.2 | 6.0 |
| NDF (% of DM) | — | 22.1 | 46.1 |
| ADF (% of DM) | – | 10.6 | 28.9 |
| Gross Energy (Mcal/kg of DM) | 4.5 | 4.3 | 4.5 |

<sup>1</sup> Prewaning diet management = 5 L of MR (15% DM) + free access to starter.
Postweaning management = 3 kg starter + free access to corn silage.
<sup>2</sup>Powder integral milk, wheat isolated protein, acidifying additive, whey, coconut oil, palm oil, vitamin A, Vitamin D3, Vitamin E, Vitamin C.
<sup>3</sup>Basic composition: oats (rolled grains), calcitic limestone, sodium chloride, corn gluten meal, defatted corn germ, wheat bran, soybean meal, rice hulls, kaolin, molasses, flocculated corn, ground corn, corn grain, alfalfa hay, monensin, citrus pulp, dried sugarcane yeast, whole toasted soybean, sodium selenite, copper sulfate, manganese sulfate, cobalt sulfate, iron sulfate, zinc sulfate, calcium iodate, vitamin A, vitamin B1, vitamin B12, vitamin B2, vitamin B6, vitamin C, vitamin D3, vitamin E, vitamin K, niacin, pantolenic acid, folic acid, biotin, propionic acid, caramel aroma, milk aroma and probiotic additive.

At 4 d of age, heifers started to be fed with 5 L/d of a MR (Kalvolak, Nutrifeed, Netherlands; Table 1) reconstituted at 15% (dry matter basis), divided into two equal meals (0800 and 1600 h) in buckets provided with rubber teats (Milkbar, New Zealand). The heifers were allocated to one of the two treatments with 4 d of age, maintaining a balance of birth month, birth BW, genetic composition, and total serum protein: Control no additive (CON; n = 15) and blend of essential oils supplementation (BEO, 1 g/d/calf, Apex Calf, Adisseo, China; n = 14). Apex Calf is a dry powder that contains a blend of plant extracts derived from anise, cinnamon, garlic, rosemary and thyme. The blend of essential oil of each meal was weighed to have 0.5 g and kept in 15 ml tubes in a dark box. They were then mixed with a 10 mL of milk, homogenized and incorporated in 0.49 L of MR (0.5 g/calf at morning meal and 0.5 g/calf at afternoon meal) to ensure total ingestion of the product. After ingesting 0.5 L MR with 0.5 g of the blend of essential oils, the rest of the meal was given.

Heifers were weaned at 60 d of age. During the postweaning period, from 61 to 90 d of age, all heifers received starter and corn silage (Table 1). The amount of corn silage provided was enough to result in at least in 10 % orts, and the starter intake was fixed in maximum 3.0 kg calf/d, divided into two meals. All heifers were dehorned at 70 d of age and received local anesthesia (5.0 mL/horn, Lidovet, Bravet, Engenho Novo, Brazil) and 2 d of non-steroid anti-inflammatory (0.025 mL/kg, Maxicam 2%, Ouro fino, Cravinhos, Brazil).

### Intake and nutritional composition analysis

Feed intake (MR, starter, water and corn silage) was measured daily. Samples of MR, starter, and corn silage were collected three times a week to obtain a week pool for nutritional analyses. Samples of starter and corn silage were oven dried at 55 °C for 72 h and ground in Wiley mill (model 3, Arthur H. Thomas Co., Philadelphia, PA) through a 1-mm screen before analysis. Starter, corn silage and MR were analyzed to determine DM (Method 934.01), CP (Method 988.05), ether extract (Method 920.39), ash (Method 942.05), according to AOAC (18). NDF and ADF were determined in sequence using the method described by Van Soest et al. (19). Gross energy was determined using an adiabatic bomb calorimeter (Parr Instrument Company, Moline, IL).

### Structural growth

BW was measured on birth date, 3 d of age, and, thereafter, every 3 d before the morning meal using a weighing-machine (ICS 300, Coimma, Dracena, Brazil). Wither height (distance from the base of the front feet to the withers), rump height (distance from the base of the rear feet to the rump), rump width (distance between ileus) and heart girth (circumference of the chest) were measured on birth date and, thereafter, every 7 d until the end of the experiment. These measurements were taken on a flat surface using a portable hypometer and a measuring tape. Feed efficiency was calculated using ADG and DMI ratio (20).

### Rumen fermentation

Rumen fluid samples were collected through an oroesophageal tube 4 h after morning feeding at 14, 28, 42, 60, 74 and 90 d of age and pH was assessed using a portable potentiometer (Phmetro T-1000, Tekna, Araucária, Brazil). Ruminal fluid (10 mL) was acidified with 1 mL of 20% metaphosphoric acid, and 2 mL of 50% sulfuric acid for VFA and nitrogen ammonia analyses. These samples were stored at -20 °C for further analysis. Nitrogen ammonia concentration was quantified using the colorimetric distillation method proposed by Chaney and Marbach (21), and its absorbance was measured at 630 nm (Termo Fisher Scientific, Madison, USA) after Kjedahl distillation with magnesium oxide and calcium choride according to Method 920.03 (18). VFA concentrations were determined in the samples previously centrifuged at 1,800 × *g* for 10 min at room temperature (22 – 25 ° C) by high performance liquid chromatography (Waters Alliance e2695 Chromatograph, Waters Technologies do Brazil LTDA, Barueri, SP, Brazil).

### Blood cell count, metabolites and IGF-1

Jugular blood samples were collected from all animals at birth before colostrum feed and, thereafter, on days 4, 7, 14, 21, 28, 35, 42, 49, 56, 63, 74, 81 and 90, three hours after morning feeding, for beta-hydroxybutyric acid (BHB), urea and glucose and, on days 4, 14, 28, 42, 56, 63, 74, 81 and 90 d, for IGF-1 concentrations. Blood samples were collected into tubes without anticoagulant (for BHB and urea), with sodium fluoride (for glucose) or with heparin for IGF-1, (Labor Import, Osasco, Brazil). They were immediately transported on ice to the laboratory and were centrifuged at 3000 x *g* for 10 min at room temperature (22 – 25 °C). Two aliquots of each metabolite and hormone sample were individually allocated into microtubes and frozen at -20 °C for further analysis. BHB and urea serum concentrations were determined by use of an auto-analyzer (Cobas Mira Plus, Roche Diagnostic Systems, Risch-Rotkreuz, Switzeland) using commercial kits (Ranbut-D-3-Hidroxibutyrate, Randox Laboratories Ltd., Antrim, UK; Urea UV, Kovalent do Brasil Ltda., Bom Retiro São Gonçalo, Brazil). Plasma glucose was measured in a microplate Spectrophotometer EON (Biotek Instruments Inc., Winooski, VT) using the enzymatic colorimetric method (Kovalent do Brasil Ltda., Rio de Janeiro, Brazil). IGF-1 plasma concentrations were analyzed using chemiluminescence essay (Immulite2000 Systems 1038144, IGF-1 200, Siemens Healthcare Diagnostics Products Ltd., Llanberis, Gwynedd, UK).

Blood samples were collected for complete blood count during preweaning at 0, 30 and 60 d of age, by jugular vein puncture into EDTA tubes (Labor Import, Osasco, Brazil), and immediately transported on ice to the laboratory. An automatic hematology cell counter (SDH – 3 vet, Labtest Diagnóstica S.A., Brazil) was used to perform: red blood cell count (RBC), packed cell volume (PCV), hemoglobin (Hb), mean corpuscular volume (MCV), mean corpuscular hemoglobin concentration (MCHC), platelet and total white blood cell count. Manual white cell blood differential counting was also performed by microscopic examination evaluating 100 leukocytes in a 1,000 x microscopic magnification for: total leukocyte count, basophils, eosinophils, neutrophils, band neutrophils, segmented neutrophils, lymphocytes, monocytes. Morphological changes were also performed, such as toxic neutrophils, reactive lymphocytes and activated monocytes (22). With these results it was also performed platelet to lymphocytes ratio (PLR) and neutrophils to lymphocytes ratio (NLR).

### Health measurements

Health measurements (fecal and respiratory scores) were performed daily, in the morning, before other animal management. Fecal scores were graded according to McGuirk (2), as follows: 0 – normal (firm but not hard); 1 – soft (does not hold form, piles but spreads slightly); 2 – runny (spreads readily to about 6 mm depth); and 3 – watery (liquid consistency, splatters). A heifer was considered to have diarrhea if fecal score was 2 or 3. Severe diarrhea was considered severe when fecal score was 3.

Daily respiratory score evaluations were adapted from the University of Wisconsin calf health scoring chart (2), considering rectal temperature score: 0 – temperature between 37.8 and 38.3 °C, 1 – temperature between 38.4 and 38.8 °C, 2 – temperature between 38,9 and 39.3 °C, 3 – temperature above 39.4 °C; cough score: 0 – none, 1 – induce single cough, 2 – induced repeated or occasional spontaneous coughs, 3 – repeated spontaneous coughs; nose score: 0 – normal serous discharge, 1 – small amount of unilateral cloudy discharge, 2 – bilateral cloudy or excessive mucus discharge, 3 – copious bilateral mucopurulent discharge; eye score: 0 – normal, no discharge, 1 – small amount of ocular discharge, 2 – moderate amount of bilateral discharge, 3 – heavy ocular discharge; ear score: 0 –normal, 1 – ear flick or head shake, 2 – slight unilateral drop, 3 – head tilt or bilateral drop. Final respiratory score considered the sum of all punctuations.

Heifers were treated with anti-inflammatories only if respiratory score sum was above 4 or if they presented fever for two days subsequently. Fever was considered when pre-meal morning temperature was ≥ 39.4 °C. Antibiotics were used only when pulmonary commitment was detected, or fever combined with diarrhea for 2 d subsequently.

### Minimum inhibitory concentration

Broth dilution method was used to evaluate the minimum inhibitory concentration (MIC) of BEO against two relevant enteric bacteria: enterotoxigenic *Escherichia coli* (K99^+^ strain) and *Salmonella typhimurium* previously isolated from an outbreak in calves (Ramos et al., 2019). Two different preparations of BEO product were used to perform MIC: a - homogenized in purified water; b - homogenized in a solution with 3.0 g of isopropyl myristate, 8.25 g of propylene glycol, 7.25 g of tween 80 and 100 ml of water. Both preparations were submitted to 0.22 µm filtration. A solution with an initial concentration of 1.0 µg/mL was submitted to serial dilutions from 1:2 to 1:256 in 96-wells plates. Thus, 100 µL of a solution containing 5 x 10^5^ CFU/mL of the two selected bacteria. After overnight incubation at 35 °C, microtiter plates were examined for visible bacterial growth evidenced by turbidity and color change.

### Statistical analysis

Statistical analysis was conducted utilizing R^®^ (R Core Team, 2019). The data collected was summarized by period (pre-weaning – 4 to 60 d and post-weaning – 61 to 90 d) and per week within each period. A randomized block experimental design with repeated measures was implemented to test the hypothesis of the effect of the blend of essential oils on each performance outcome. More specifically, the outcomes analyzed were feed intake, structural growth, ruminal, blood and health parameters. The control group was assigned 15 experimental units (CON), while the treatment was assigned 14 (BEO).

The analysis of each outcome was performed independently of all others using linear mixed models (package: nlme). Each independent outcome was modeled as a function of the following fixed effects: treatment, experimental week, the interaction between treatment and week. Birth weight and serum Brix refractometer were tested as a covariate but did not improve statistical significance. Therefore they were eliminated from the model. The genetic composition of the animal was included as a blocking effect. The effect of heifer within treatment was included in the models to account for individual variability.

The continuous outcomes such as intakes, performance, ruminal and blood parameters were analyzed with ANOVA. A 95% Confidence Interval was adopted to accept or deny the null hypothesis and P-values were produced with a Fisher test. In order to meet the required assumptions of this model, all outcomes were tested for normality, and variable transformation was applied to milk replacer intakes to meet that assumption.

The categorical outcomes fecal and respiratory scores were analyzed using a non-parametric aligned rank transformation test, implemented in the R package ARTool. A 95% Confidence Interval was also adopted for the non-parametric tests.

## Results and discussion

### Intake and heifer performance

Most studies evaluating essential oils or a supplement with BEO to dairy calves, feed the additive in the starter, to benefit rumen development and accelerate growth. However, the intake of starter in the firsts weeks of age is small (23). Due to the calf’s limited capability of ingesting large solid feed amounts in the first days of life, the supplement intake could be compromised during pre-weaning period and possibly mask any effects. Therefore, in this trial, it was decided to offer BEO in the liquid diet.

The tested BEO has a strong aroma. Nevertheless, due to the way it was offered, no rejection of the mixture BEO and MR was observed. The supplemented heifers consumed liquid diet equally to the control treatment, with no refusal and good acceptance (Table 2). Some essential oils have different acceptability by animals. Chapman et al. (24) testing cinnamaldehyde essential oil in weaned dairy heifers fed a total mix ratio, observed that the animals preferred the taste in the control treatment, and the modification of feed intake was related to problems of palatability with the essential oil used in the experiment. Differences between flavor and palatability of BEOs could be due to the way of supply, as well as to essential oil plant source.

**Table 2.** Pre and post-weaning milk replacer (MR) intake, starter intake, total dry matter intake (DM), total crude protein intake (CP), total gross energy and water intake of heifers (n = 29) of control (CON) and supplemented with blend essential oils (BEO) in milk replacer during pre-weaning.

| Intake | Treatment |  | SEM | P – value<sup>3</sup> |  |  |
| --- | --- | --- | --- | --- | --- | --- |
|  | CON<sup>1</sup> | BEO<sup>2</sup> |  | T | W | T x W |
|  | (n =15) | (n =14) |  |  |  |  |
| Pre-weaning (4 to 60 d) |  |  |  |  |  |  |
| MR (kg of DM/d) | 0.71 | 0.71 | 0.02 | 0.30 | <0.001 | <0.001 |
| Starter (kg of DM/d) | 0.30 | 0.31 | 0.02 | 0.92 | <0.001 | 0.82 |
| Total DM (kg/d) | 1.00 | 1.16 | 0.06 | 0.58 | <0.001 | 0.31 |
| Total CP (kg/d) | 0.19 | 0.18 | 0.01 | 0.58 | <0.001 | 0.31 |
| Total gross energy (Mcal/kg) | 4.51 | 4.59 | 0.12 | 0.58 | <0.001 | 0.30 |
| Water (kg/d) | 1.39 | 1.30 | 0.32 | 0.98 | <0.001 | 0.64 |
| Post-weaning (61 to 90 d) |  |  |  |  |  |  |
| Starter (kg of DM/d) | 1.84 | 2.02 | 0.28 | 0.39 | <0.001 | 0.31 |
| Corn Silage (kg of DM/d) | 0.12 | 0.11 | 0.03 | 0.51 | <0.001 | 0.26 |
| Total DM (kg/d) | 1.97 | 2.14 | 0.29 | 0.39 | <0.001 | 0.32 |
| Total CP (kg/d) | 0.44 | 0.47 | 0.07 | 0.36 | <0.001 | 0.72 |
| Total gross energy (Mcal/kg) | 8.61 | 9.35 | 1.34 | 0.39 | <0.001 | 0.29 |
| Water (kg/d) | 5.41 | 5.69 | 0.84 | 0.61 | <0.001 | 0.10 |

<sup>1</sup>CON = control; <sup>2</sup>BEO = 1 g/calf/d blend of essential oil.
<sup>3</sup>T = treatment effect; W= week effect, T x W = treatment by week interactions.

However, besides fixed MR intake amount, there was a week effect and a week and treatment interaction effect (*P ≤* 0.05, Table 2, Fig 1). At the end of week 1 until week 3, heifers had diarrhea and this event impacted on MR intake. Differences between treatments were greater in those weeks when compared with the other weeks, with lower intake for CON group. It was found an effect between fecal scores and MR intake (*P ≤* 0.001), besides low correlation value (−0.25). Thus, results reveled an inverse association between both parameters, were greater fecal scores impacted on smaller MR intake, and vice versa.

**Fig 1.**
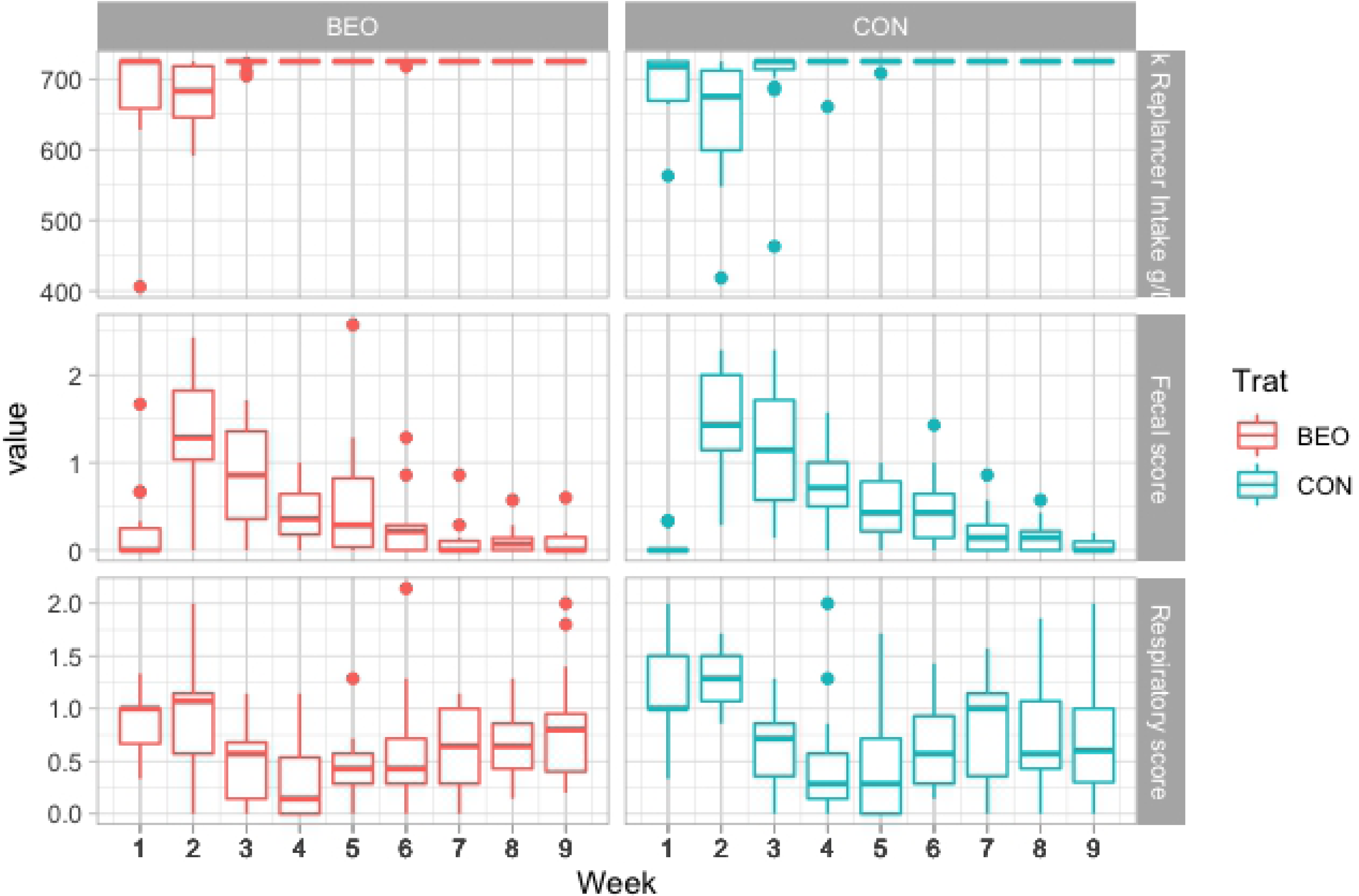
MR intake (g of DM/d), respiratory and fecal scores of control heifers (CON) and heifers supplemented with 1.0 g/calf/ d of blend of essential oils (BEO) in milk replacer during preweaning period.

Intake of starter, water, total DM, CP, and gross energy, ADG and feed efficiency were not affected by treatment during pre- and post-weaning (Table 2 and 3), indicating that the palatability and aroma of BEO did not discourage the heifers’ MR consumption nor the starter and water intakes. Santos et al. (25) tested EO supplementation for dairy calves using two routes of supplementation (MR and starter), and had similar results, for intake, BW and ADG during preweaning. However, Kazemi-Bonchenari et al. (16) and Liu et al. (15) evaluated commercial source of essential oils in starter and found better ADG and feed efficiency during the preweaning period for supplemented calves, as well as higher BW during weaning. As for carryover effect, differently from this experiment, Salazar et al. (26) observed calves with BEO treatment in starter presenting higher ADG and feed efficiency.

**Table 3.** Pre and post-weaning performance and body development of heifers (n = 29) of control (CON) and supplemented with essential oils blend (BEO) in milk replacer during pre-weaning.

| Item | Treatment |  | SEM | *P* – value<sup>3</sup> |  |  |
| --- | --- | --- | --- | --- | --- | --- |
|  | CON<sup>1</sup> | BEO<sup>2</sup> |  | T | W | T x W |
|  | (n = 15) | (n = 14) |  |  |  |  |
| **Performance** |  |  |  |  |  |  |
| Birth BW (kg) | 32.40 | 31.97 | 0.59 | 0.85 | – | – |
| Weaning BW (kg) | 64.36 | 66.66 | 1.07 | 0.45 | – | – |
| Final BW (kg) | 89.88 | 93.34 | 1.57 | 0.57 | – | – |
| ADG preweaning (kg) | 0.55 | 0.53 | 0.02 | 0.49 | <0.001 | 0.23 |
| ADG postweaning (kg) | 0.81 | 0.84 | 0.27 | 0.76 | 0.001 | 0.60 |
| Feed efficiency preweaning | 0.62 | 0.56 | 0.008 | 0.06 | <0.0001 | 0.29 |
| Feed efficiency postweaning | 0.44 | 0.42 | 0.04 | 0.50 | 0.68 | 0.42 |
| **Body measures** |  |  |  |  |  |  |
| **Prewaning (4 to 60 d)** |  |  |  |  |  |  |
| Withers height (cm) | 72.74 | 72.59 | 1.25 | 0.86 | <0.001 | 0.48 |
| Rump height (cm) | 75.89 | 75.90 | 0.66 | 0.98 | <0.001 | 0.62 |
| Rump width (cm) | 19.03 | 19.42 | 0.66 | 0.23 | <0.001 | 0.94 |
| Heart girth (cm) | 80.70 | 81.50 | 0.009 | 0.34 | <0.001 | 0.68 |
| **Postweaning (61 to 90 d)** |  |  |  |  |  |  |
| Withers height (cm) | 82.66 | 82.55 | 1.06 | 0.92 | <0.001 | 0.72 |
| Rump height (cm) | 86.02 | 86.64 | 1.07 | 0.61 | <0.001 | 0.80 |
| Rump width (cm) | 22.59 | 22.99 | 0.43 | 0.28 | <0.001 | 0.40 |
| Heart girth (cm) | 96.55 | 97.85 | 1.44 | 0.27 | <0.001 | 0.40 |

<sup>1</sup>CON = control; <sup>2</sup>BEO = 1 g/calf/d blend of essential oil.
<sup>3</sup>T = treatment effect; W= week effect, T x W = treatment by week interactions.

### Structural growth

Body development was not affected by BEO supplementation in MR (Table 3) during pre and post-weaning. As was also observed for intake and ADG, a week effect (P ≤ 0.05) was detected in all variables due to normal animal growth. Kazemi-Bonchenari et al. (16) suggested that essential oils supplementation could only be effective in structural growth when associated with higher protein concentration in the starter, since the authors found an interaction between protein level and essential oils supplementation. However, Liu et al. (15) feeding an amount of 44.1 ppm of a commercial essential oil prebiotic, did find increased body frame measurements and body weight for supplemented calves, suggesting that feeding essential oils could enhance growth performance if fed at an appropriate rate and in a determined amount.

### Rumen fermentation

There were no differences in ruminal pH for CON and BEO treatments during the pre-weaning period. During the post-weaning period the BEO treatment present lower pH (*P* ≤ 0.05, Table 4). Since there were no differences between treatments during pre-weaning, the carryover effect may not be assumed to be the answer for this difference. Although no differences on intake were observed, heifers’ ingestion behavior might justify the difference in post-weaning pH. However, this parameter was not evaluated. Nevertheless, previous studies with essential oils supplementation also found low pH values for pre-weaning calves, but no essential oils treatment effect (16,25,27).

**Table 4.** Pre and post-weaning rumen mean values of rumen pH, ammonia nitrogen (Ammonia-N) and volatile fatty acids (VFA) of control heifers (CON) and heifers supplemented with essential oils blend (BEO) in milk replacer during pre-weaning.

| Item | Treatment |  | SEM | *P* – value<sup>3</sup> |  |  |
| --- | --- | --- | --- | --- | --- | --- |
|  | CON<sup>1</sup> | BEO<sup>2</sup> |  | T | W | T x W |
|  | (n = 15) | (n = 14) |  |  |  |  |
| **Pre-weaning (4 to 60 d)** |  |  |  |  |  |  |
| Rumen pH | 5.99 | 5.85 | 0.52 | 0.37 | 0.03 | 0.06 |
| Rumen ammonia-N (mg/dL) | 11.40 | 13.80 | 0.03 | 0.15 | <0.001 | 0.37 |
| Rumen VFA (μmol/mL) |  |  |  |  |  |  |
| Acetic (C2) | 30.80 | 27.16 | 8.15 | 0.24 | <0.001 | 0.14 |
| Propionic (C3) | 18.88 | 20.01 | 7.11 | 0.59 | <0.001 | 0.14 |
| Butyric (C4) | 0.80 | 0.80 | 0.08 | 0.83 | 0.005 | 0.98 |
| C2:C3 | 1.97 | 1.69 | 0.12 | 0.05 | <0.001 | 0.95 |
| **Post-weaning (61 to 90 d)** |  |  |  |  |  |  |
| Rumen pH | 6.19 | 5.90 | 0.001 | 0.05 | 0.001 | 0.86 |
| Rumen ammonia-N (mg/dL) | 10.97 | 9.53 | 9.03 | 0.17 | 0.91 | 0.88 |
| Rumen VFA (μmol/mL) |  |  |  |  |  |  |
| Acetic (C2) | 38.32 | 39.03 | 8.48 | 0.81 | 0.006 | 0.93 |
| Propionic (C3) | 28.27 | 30.69 | 5.16 | 0.41 | 0.003 | 0.75 |
| Butyric (C4) | 5.94 | 6.16 | 1.19 | 0.82 | 0.95 | 0.62 |
| C2:C3 | 1.43 | 1.23 | 0.20 | 0.006 | 0.74 | 0.93 |

<sup>1</sup>CON = control; <sup>2</sup>BEO = 1 g/calf/d blend of essential oil.
<sup>3</sup>T = treatment effect; W= week effect, T x W = treatment by week interactions.

Low pH could indicate acute or sub-acute ruminal acidosis or interfere with animal performance. However, signs of tympanism, gas bubbles or abdominal discomfort were not observed among the animals of this experiment. Young calves have a poorly developed rumen, with an immature epithelium to absorb large amounts of VFA. Possibly, for that reason, a low pH is frequently found in pre-weaning calves. Therefore, acidosis has not been reported at this phase, since calves can stand lower rumen pH values when compared to adult ruminants (23).

Besides the previous cited effects, essential oils supplementation is related to antimicrobial and antifungal effects (9,28). Essential oils cause hydrophobicity, disrupting bacteria membrane, increasing permeability and causing in a toxic effect (7,12). This activity could result in inhibition of ruminal deamination and methanogenesis (13). As a consequence, it would result in a decrease of the ruminal nitrogen ammonia, methane and acetate concentrations, and an increase of the propionate and butyrate concentrations (28). Changes in these profiles in rumen fluid would also alter the acetate:propionate (C2:C3) proportions. Since butyrate and propionate are important for ruminal papillae development, and especially propionate is used in the gluconeogenesis route (5), a smaller C2:C3 ratio is wanted. In this experiment, BEO supplementation reduced the C2:C3 proportion during the pre- (*P* ≤ 0.05) and postweaning phases (*P* ≤ 0.05) (Table 4). Confirming these findings, Kazemi-Bonchenari et al. (16) registered lower C2:C3 proportion for calves in both groups supplemented with essential oils in starter compared with non-essential oils groups (1.56 and 1.47 – group without essential oils supplementation x 2.02 and 1.77 – group with essential oils supplementation; *P <* 0.01). However, in our experiment, essential oils were provided mixed in small amounts of MR to ensure the whole intake of the product. This could reduce the chances of essential oils entering the rumen and alter local microbiota as well as VFA. Nevertheless, since the MR amount was small and given at beginning of the feeding, it is possible that the esophageal groove was still open, permitting essential oils content extravasation to the rumen. Akbarian-Tefaghi et al. (27) also found higher C2:C3 proportion during the preweaning phase for calves suplemented with thyme essential oils when compared with control the group (2.25 x 1.78).

Despite this, in this study, VFA concentrations were not affected by BEO supplementation during pre- and post-weaning, but increased with age (*P* ≤ 0.05, Table 4). Corroborating these findings, Santos et al. (25) did not present differences for VFA in supplemented calves, but also observed age impact on the VFA profile. Since VFA are associated with rumen development, a high starter intake at older age could result in increase in the VFA parameters, explaining age impact.

As for nitrogen ammonia, there was no treatment effect during pre- or postweaning. In contrast, Santos et al. (25) reported greater nitrogen ammonia, suggesting that essential oils could not modulate deamination nor the population of ammonia producing bacteria.

For all ruminal parameters it was found week effect during preweaning (*P* ≤ 0.05, Table 4). That could be related to the increase in starter intake, rumen development and microbiota colonization, since weeks 3 and 5 parameters were different from week 7 and 9 parameters.

### Blood cell count, metabolites and IGF-1

All blood metabolites, during pre- and postweaning period, were not altered by BEO supplementation (Table 5). Similar patterns of BHB, glucose (14,25), urea (16), total plasma protein, and IGF-1 (29) were found for essential oil supplemented calves. Nevertheless, BHB and urea increased with age (*P* ≤ 0.05, Table 5), since they are directly correlated with fatty acid metabolism and ruminal ammonia concentration, respectively (30). IGF-1 increased with age on the preweaning phase (*P* ≤ 0.05). Since this hormone is a mitogen and related with cell proliferation and differentiation, it is correlated with BW and animal growth (31).

**Table 5.** Pre and post-weaning mean blood concentrations of insulin growth factor type 1 (IGF-1) and metabolites of control heifers (CON) and heifers supplemented with blend of essential oils blend (BEO) in milk replacer during pre-weaning.

| Item | Treatment |  | SEM | P – value<sup>3</sup> |  |  |
| --- | --- | --- | --- | --- | --- | --- |
|  | CON<sup>1</sup> | BEO<sup>2</sup> |  | T | W | T x W |
|  | (n = 15) | (n =14) |  |  |  |  |
| Pre-weaning (4 to 60 d) |  |  |  |  |  |  |
| BHB (mmol/L) | 0.17 | 0.12 | 0.02 | 0.43 | 0.001 | 0.98 |
| Urea (mg/dL) | 24.55 | 22.69 | 3.76 | 0.16 | 0.02 | 0.31 |
| Glucose (mg/dL) | 100.35 | 102.97 | 16.50 | 0.49 | 0.15 | 0.56 |
| IGF-1 (ng/mL) | 101.95 | 93.16 | 32.4 | 0.38 | <0.001 | 0.27 |
| Post-weaning (61 to 90 d) |  |  |  |  |  |  |
| BHB (mmol/L) | 0.36 | 0.37 | 0.10 | 0.70 | <0.001 | 0.13 |
| Urea (mg/dL) | 24.57 | 22.73 | 4.34 | 0.16 | 0.01 | 0.34 |
| Glucose (mg/dL) | 88.45 | 84.74 | 8.65 | 0.29 | 0.22 | 0.14 |
| IGF-1 (ng/mL) | 160.70 | 175.94 | 23.4 | 0.43 | 0.31 | 0.12 |

<sup>1</sup>CON = control; <sup>2</sup>BEO = 1 g/calf/d blend of essential oil.
<sup>3</sup>T = treatment effect; W= week effect, T x W = treatment by week interactions.

Glucose did not change during pre-weaning phase and decreased during the post-weaning period (Table 5). Taking into accounts that calves use glucose as a primary source of energy in the firsts weeks of age, these age related changes are associated with changes in diet and rumen development (32). After weaning, calves complete their rumen development and VFA produced by ruminal microbiota become the main energy source, justifying BHB concentration increase and glucose concentration decrease (5,33). However, since there were changes in C2:C3 proportion in BEO group, the increase of propionic acid could consequently impact on glucose blood concentration. Since essential oils can increase of insulin sensitivity, not finding glucose differences between treatments does not mean that there were no changes in glucose pathway. Therefore, further investigations over theses aspects is needed.

All blood cells count were within age and specie normality. According to Benesi et al. (34) changes in blood cells count are normal during heifer growth, and blood cells tend to increase with animal age which corroborated our findings of week effect on mean corpuscular volume (MCV), basophils, eosinophils, segmented neutrophils, lymphocytes, monocytes and platelets (*P* ≤ 0.05). There were no differences in erythrogram parameters between BEO and CON (Table 6). Leukogram parameters showed decreased counts of basophil and platelet cells in BEO treatment (*P* ≤ 0.05). An interesting interaction effect was found also for lymphocytes (Fig 2), where values of d 30 and 60 were different from d 1, what was expected since cells changes with animal growth. Basophils and platelets are originated from different myeloid precursors and both play important roles on inflammation balance and immune response development in mammal (35). The lower counts of basophil and platelets on BEO treatment may influence and modulate inflammatory response by secretion of immune modulators (36), growth factors or chemotaxis on variety of white blood cells (22).

**Table 6.** Pre-weaning hematological parameters of control heifers (CON) and heifers supplemented with blend of essential oils blend (BEO) in milk replacer during pre-weaning.

| Item<sup>1</sup> | Treatment |  | SEM | *P* – value<sup>4</sup> |  |  |
| --- | --- | --- | --- | --- | --- | --- |
|  | CON<sup>2</sup> | BEO<sup>3</sup> |  | T | W | T x W |
|  | (n =15) | (n =14) |  |  |  |  |
| RBC (x 10<sup>6</sup>/μL) | 8.02 | 7.95 | 0.88 | 0.86 | 0.63 | 0.87 |
| PCV (%) | 35.53 | 35.21 | 5.05 | 0.85 | 0.11 | 0.69 |
| Hb (g/dL) | 11.07 | 10.94 | 1.61 | 0.81 | 0.14 | 0.73 |
| MCV (fL) | 44.74 | 44.51 | 2.94 | 0.74 | <0.001 | 0.51 |
| MCHC (%) | 31.10 | 31.14 | 0.76 | 0.87 | 0.15 | 0.99 |
| Total leukocytes<br>(/μL) | 10,908.45 | 11,200.78 | 2,630.0 | 0.76 | 0.19 | 0.22 |
| Basophils (/μL) | 2.14 | 0.00 | 1.03 | <0.001 | <0.001 | <0.001 |
| Eosinophils (/μL) | 68.40 | 143.90 | 0.66 | 0.24 | <0.001 | 0.36 |
| Band neutrophil<br>(/μL) | 31.76 | 26.22 | 5.69 | 0.68 | 0.83 | 0.31 |
| Segmented<br>neutrophils (/μL) | 5,300.63 | 5,286.56 | 1,700.0 | 0.98 | <0.001 | 0.78 |
| Lymphocytes (/μL) | 4,837.40 | 5,082.82 | 1,120.0 | 0.66 | <0.001 | 0.01 |
| Monocytes (/μL) | 421.60 | 466.00 | 247.0 | 0.48 | 0.01 | 0.29 |
| Platelet (x 10<sup>3</sup>/μL) | 410.41 | 353.70 | 108.0 | 0.04 | <0.001 | 0.10 |
| Plasmatic protein<br>(g/dL) | 6.03 | 6.03 | 0.72 | 1.00 | 0.17 | 0.40 |
| PLR | 0.08 | 0.08 | 0.03 | 0.91 | 0.02 | 0.04 |
| NLR | 1.26 | 1.46 | 0.03 | 0.60 | <0.001 | 0.55 |

<sup>1</sup>RBC: red blood cell, PCV: packed cell volume, Hb: hemoglobin, MCV: mean corpuscular volume, MCHC: mean corpuscular hemoglobin concentration, PLR: platelet lymphocyte ratio, NLR: neutrophils lymphocytes ratio.
<sup>2</sup>CON = control; <sup>3</sup>BEO = 1 g/calf/d blend of essential oil.
<sup>4</sup>T = treatment effect; W= week effect; T x W = treatment by week interactions.

**Fig 2.**
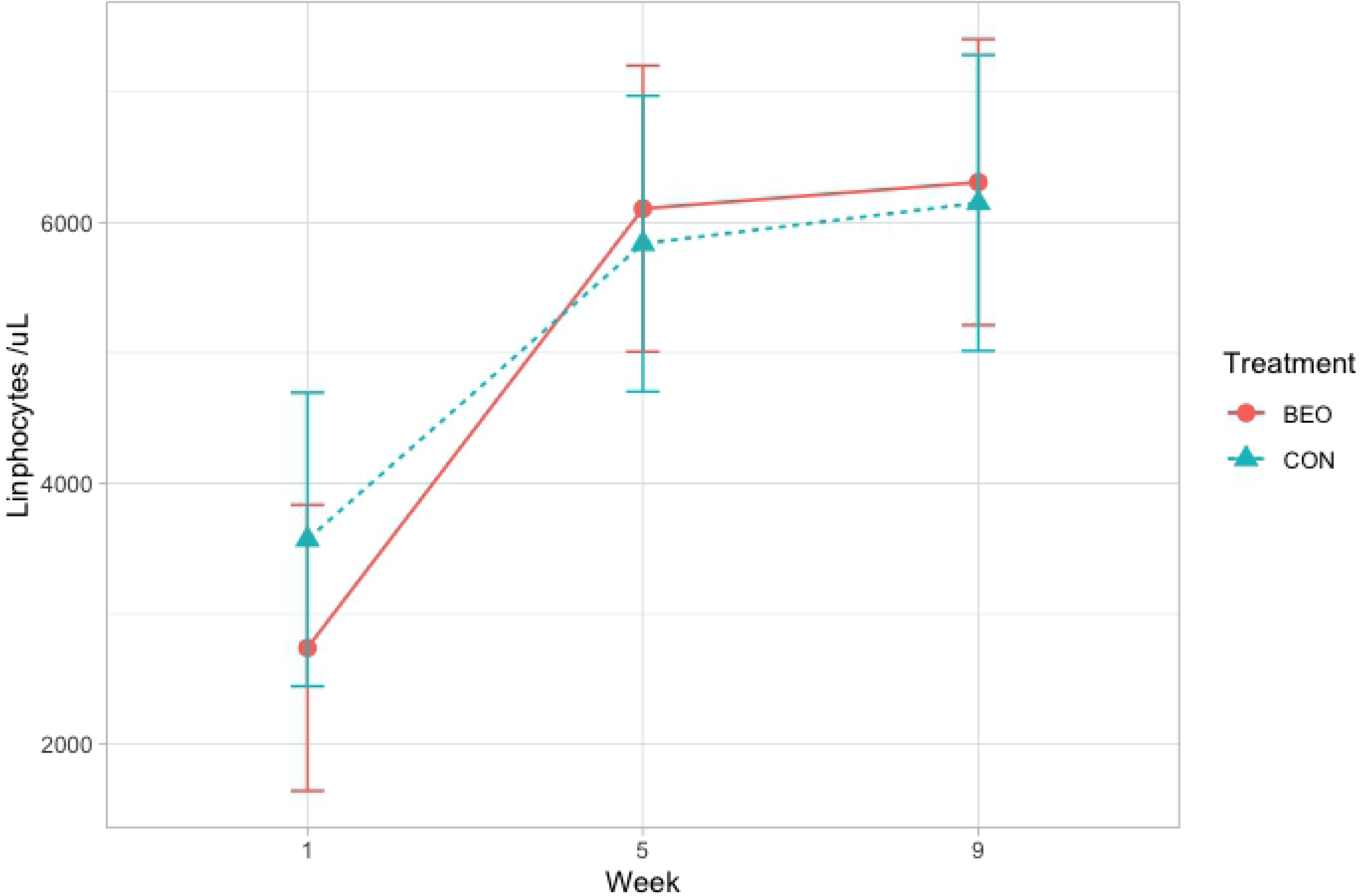
Lymphocytes values of control heifers (CON) and heifers supplemented with 1.0 g/calf/ d of blend of essential oils (BEO) in milk replacer during preweaning period.

Seirafy and Sobhanirad (37), testing oregano and thyme oils in Holstein calves, found positive differences for erythrogram parameters, lymphocytes, neutrophils and band neutrophils with higher values for treated calves. For older animals, Gabbi et al. (38) also found a linear increase in the values for lymphocyte and monocyte counts for heifers supplemented with plant extract containing essential oils. Hence, agents with antioxidant activity, like essential oils, can reduce platelet activation and consequently reduce oxidative stress and inflammation (39). Platelets also play a central role on coagulation process. Different essential oils have been used for thrombosis treatment in human acting on platelet aggregation and its thromboxane synthesis (40). Although our results demonstrate a decrease on basophil and platelet counts, it is necessary to perform novel experiments in order to characterize the effects of BEO on inflammatory and coagulation process in heifers. Differences for PLR and NLR were not found (Table 7). These ratios are inflammatory markers and inform disease activity, being a useful tool to understand inflammation pathophysiology and immune response (41).

**Table 7.**
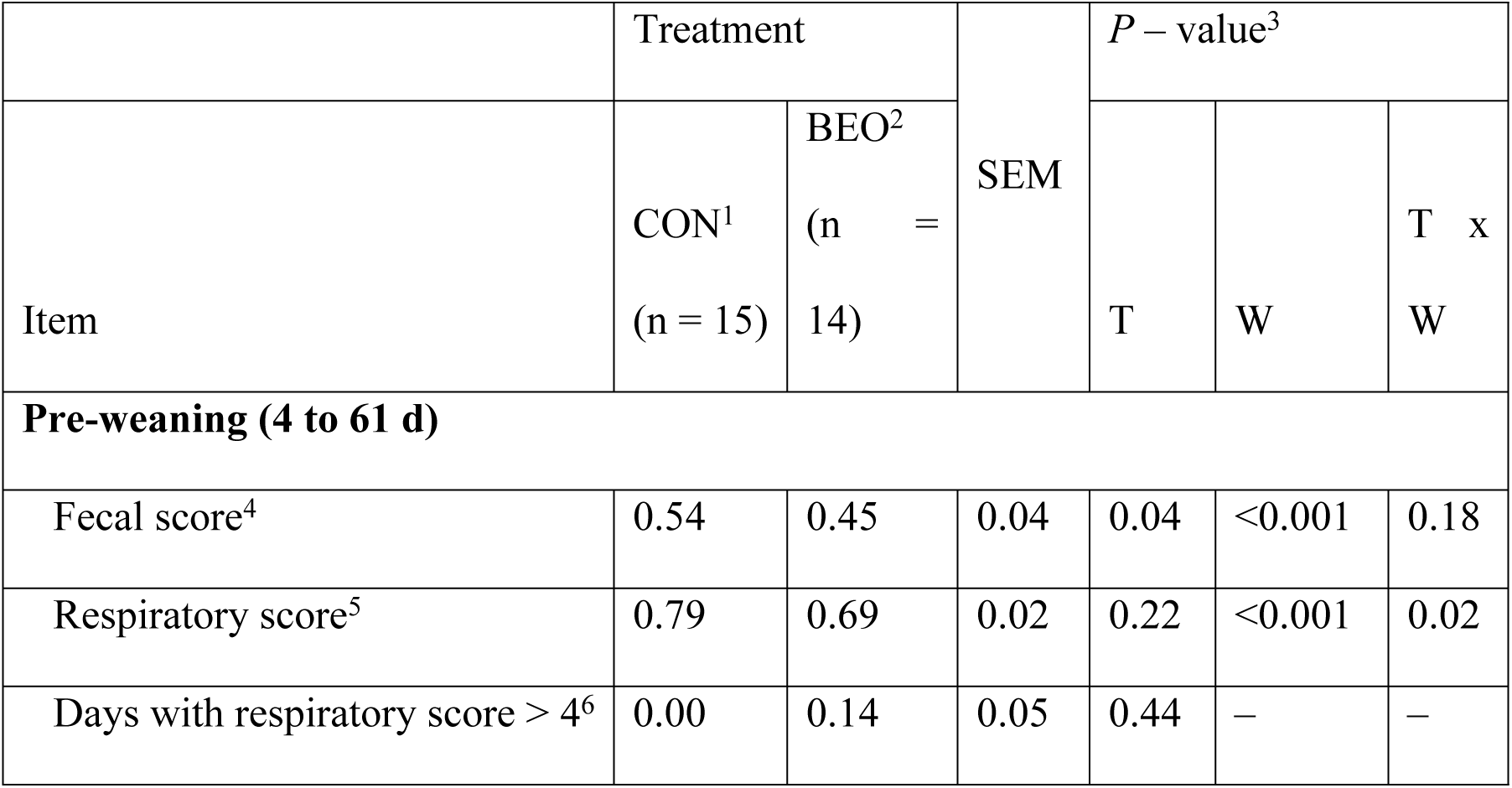
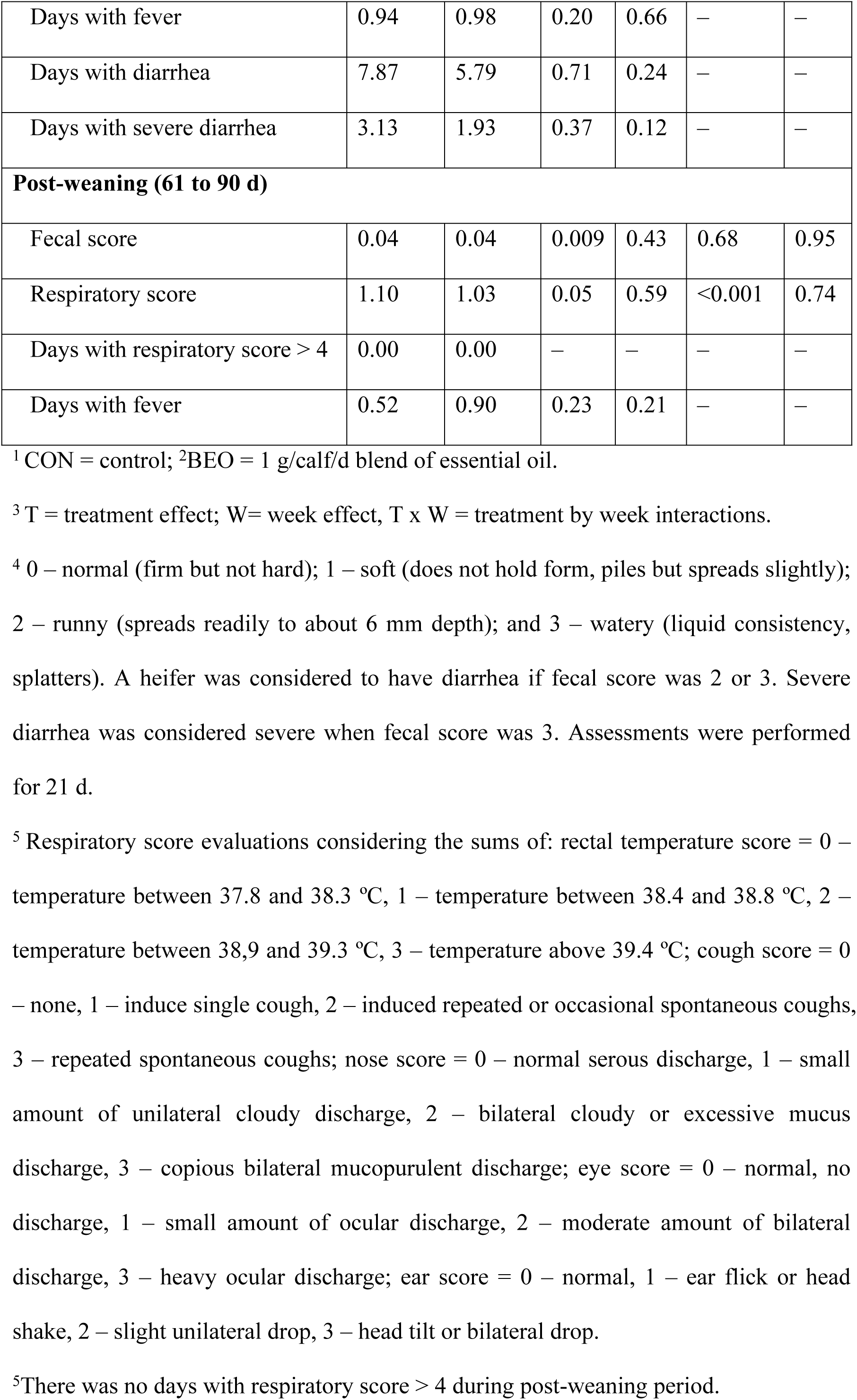
Pre and post-weaning mean values of fecal score, health score, days with health score above 4, days with fever, days with diarrhea, days with severe diarrhea of control heifers (CON) and heifers supplemented with blend of essential oils (BEO) in milk replacer during pre-weaning.

Liu et al. (15) found higher values for blood total serum protein, IgG, IgA and IgM for calves supplemented with essentials oils on the starter. Therefore, their results and the results of this trial support the theory that essential oils enhance immune status both in cellular and humoral responses.

### Health measurements and minimum inhibitory concentration

Diarrhea is the most prevalent disease for calves under one month of age. Causes for juvenile diarrhea include a combination of factors but are normally related to viral, bacterial or/and protozoa infection. Coronavirus, rotavirus, *Salmonella* spp. and *Cryptosporidium parvum* are the most common agents under 14 d of age. *Salmonella* spp., *Eimeria* spp. and *Giardia* spp. are the most common pathogens in older calves (2,42).

The average age for diarrhea occurrence was 12.2 ± 3.6 d for BEO and 13.6 ± 3.8 d for CON with no statistical difference. Diarrhea incidence on pre-weaning in BEO treatment was 85% against 93% for CON treatment with no statistical difference. Fecal score was different between treatments (*P* ≤ 0.05), with lower values for BEO, and changed through time (*P* ≤ 0.05, Table 7). Days with diarrhea and days with severe diarrhea were not different within treatments (Table 7). Three animals of each group were treated for diarrhea with anti-inflammatories, and the treatment duration was of 1.6 ± 0.57 d of treatment for BEO and 3.0 ± 1 d for CON. It is noteworthy that this treatment was done outside the total cell count.

Evaluation of the respiratory score parameters indicated that 2 BEO animals and 1 of CON animals exceeded score 4, indicating respiratory disease on pre-weaning. The average days with high score was 1.0 ± 0 d for BEO. No effect was found on days with high respiratory score or number of affected animals. However, it was observed a week and an interaction week x treatment effect on pre-weaning, with difference between treatments scores and lower values for BEO group in week 2 (*P* ≤ 0.05, Table 7, Fig 1). Week 2 was the period in which animals had higher incidence of diarrhea. Thus, it can be assumed that respiratory signs were related to previous enteric disease since a higher incidence of respiratory parameters occurred after diarrhea cases. Weeks 5 and 6 showed lower score differeces between treatments as well as lower incidence of respiratory signs. The number of treated animals was 2 for BEO only during the preweaning period, with an average of treatment days of 1.3 ± 1.4, and 3 for CON with an average of treatment days of 2.0 ± 0.57. Treatments occurred only in the pre-weaning period using antibiotics and anti-inflammatories.

Pneumonia is normally associated to the post-weaning phase. However, it may affect younger calves to have it (2). Post-weaning respiratory scores revealed higher mean values, but no animals had scores above 4. There was a week effect (*P* ≤ 0.05), in week 12, probably due to weaning and dehorning stress.

It has been reported that essential oils have an antiseptic and antimicrobial activity that may be beneficial in balancing intestinal microbiota (43). Gram positive bacteria are the most sensitive to the essential oils microbial activity (7,12), but it is possible that gram negative bacteria and some types of parasites are also susceptible (9) to different essential oils. Thus, essential oils could reduce the incidence and severity of diarrhea syndrome in calves trough inhibition of coliform overgrowth (44). BEO in 1.0 µg/mL concentration did not inhibit bacterial growth – both *E. coli* and *S. tryphimurium*. Thus, at this concentration BEO did not have any antibacterial in vitro effect. However, besides no direct influence found over the bacterial evaluation, BEO calves presented differences on basophil (Table 6) and lymphocyte cell populations (Fig 2) which could be associated to modulation of inflammatory immune response. Thus, outcomes found on fecal and respiratory scores could be associated to indirect changes in hemato-biochemical parameters since essential oils have antioxidant and anti-inflammatory effect.

## Conclusions

Feeding BEO to pre-weaned heifers on MR did not affect intake, performance parameters, blood metabolites or IGF-1 concentration. However, it changed C2:C3 proportion during pre- and post-weaning periods, as well as showed signs of immunity improvement and lower fecal scores in the pre-weaning phase. Further research is needed to define the best rout and dosage as well as understand the essential oils contribution on decrease morbidity of neonatal diseases. Therefore, essential oils are a health additive option to modern production sytems, and could be used as an alternative to improve calf health and performance.

## Acknowledments

The authors thank Professor Armando Cunha Jr. for helping perform MIC analyzes, Professor Ângela Quintão, Professor Fabiola Paes Leme and Vera Carsoso Ferreira Aiken for helping on this paper. We also thank Coordenação de Aperfeiçoamento de Pessoal de Nível Superior (CAPES, Brasília, Brazil), Fundação de Amparo à Pesquisa do Estado de Minas Gerais (FAPEMIG, Minas Gerais, Brazil), Conselho Nacional de Desenvolvimento Científico e Tecnológico (CNPq, Brasília, Brazil), Instituto Nacional de Ciência e Tecnologia Ciência Animal (INCT, Viçosa, Brazil), Embrapa Dairy Cattle (Minas Gerais, Brazil) and Adisseo Company for financial support of this research.

## Author contribution

**Conceptualization:** Sandra G. Coelho, Mariana M. Campos.

**Data curation:** Joana P. Campolina, Anna Luiza Belli.

**Formal analysis:** Joana P. Campolina.

**Funding acquisition**: Sandra G. Coelho, Mariana M. Campos.

**Investigation:** Joana P. Campolina, Anna Luiza Belli.

**Methodology:** Joana P. Campolina, Sandra G. Coelho, Anna Luiza Belli and Mariana M. Campos.

**Project administration:** Sandra G. Coelho, Mariana M. Campos.

**Resources:** Sandra G. Coelho, Mariana M. Campos, Fernanda S. Machado, Luiz G. R. Pereira, Thierry R. Tomich.

**Software:** Joana P Campolina.

**Supervision:** Sandra G. Coelho, Mariana M. Campos, Fernanda S. Machado, Luiz G. R. Pereira, Thierry R. Tomich.

**Validation:** Sandra G. Coelho, Mariana M. Campos.

**Visualization**: Sandra G. Coelho.

**Writing – original draft:** Joana P. Campolina.

**Writing – review & editing:** Joana P. Campolina, Sandra G. Coelho, Anna Luiza Belli, Fernanda S. Machado, Luiz G. R. Pereira, Thierry R. Tomich, Wanessa A. Carvalho, Rodrigo O. S. Silva, Alessandra L. Voorsluys, David V. Jacob and Mariana M. Campos.

